# Direct measurement of mavacamten and deoxyATP perturbation of the SRX/DRX ratio in porcine cardiac myofibrils using a simple, accessible and multiplexed approach

**DOI:** 10.1101/2025.07.23.666147

**Authors:** Matvey Pilagov, Ateeqa Naim, Kenneth S. Campbell, Thomas Kampourakis, Michael A. Geeves, Neil M. Kad

## Abstract

Cardiac muscle adapts to varying physiological demands by modulating the number of active myosin II motors available for contraction. These motors are organized into thick filaments in the sarcomere and generate force through an ATP-dependent interaction with thin filaments that contain actin. To conserve energy, when demand is low myosin can occupy the super-relaxed (SRX) state, which acts as a reserve. Here, we build upon the earlier studies from the Cooke lab to quantify the size of this cardiac reserve using fluorescence imaging of Cy3-ATP directly in myofibrils. This approach employs a pulse-chase method and exploits the high permeability of isolated myofibrils to monitor nucleotide release *in situ*. By preserving sarcomeric architecture while enabling rapid reagent exchange, this method bridges the gap between complex single-molecule imaging and traditional stopped-flow bulk assays using MANT-ATP. Using this approach we have studied biochemical perturbation of the SRX reserve with deoxyATP and mavacamten. DeoxyATP caused large depletion of the cardiac reserve, and mavacamten increased its size, consistent with its clinical application. Our results demonstrate the utility of this technique and the potential for further enhancement using multiplexing, holding promise for future applications in health and disease.

## Introduction

Cardiac muscle continuously contracts and relaxes over the lifetime of all higher animals. It is organised into sarcomeres comprised of lateral arrays of thick filaments each one with ~300 myosin motors capable of generating force. However, the number of motors available for cardiac contraction is modulated according to the prevailing conditions, thus optimizing energy usage. Examples of dynamic changes requiring greater numbers of available myosins include β-adrenergic stimulation or raised afterload due to elevated systemic blood pressure. These regulated motors form a reserve pool of myosins and the molecular mechanisms that lead to this regulation are currently under intense investigation.

Controlling the number of available myosins allows the heart to adapt to inotropic stresses by recruiting more myosins to pump harder and faster, or when demands are low, to rest myosins. An early and important observation is that myosin heads in muscle consume ATP ~five-times slower than myosin heads in solution (Ferenczi et al., 1978). This suggests that the sarcomeric architecture may play a role in controlling energy consumption (Starr and Offer, 1978; Cooke, 2011). In this study, we offer a method to rapidly and directly estimate the amounts of active myosins in the near-native environment of muscle myofibrils.

Contraction in muscle occurs in myofibrils, long linear arrays of sarcomeres, and is generated by the double-headed motor myosin II, which forms helically arrayed ‘thick’ filaments packed hexagonally in three dimensions. Between every three thick filaments is a thin filament that is formed from actin and decorated with tropomyosin and troponin, which control access of myosin to actin (Gordon et al., 2000). Access is gated by calcium binding to the control protein troponin (subunit C), which leads to azimuthal movement of tropomyosin across the thin filament to expose myosin binding sites on actin (McKillop and Geeves, 1993; Poole et al., 2006); and is cooperatively propagated along the thin filament (Fraser and Marston, 1995; Conibear and Geeves, 1998; Desai et al., 2015). When myosin hydrolyses ATP to ADP.Pi it can then bind to actin, which stimulates the release of Pi and ADP coupled to the myosin working stroke, leading to force generation and muscle shortening. Binding of a subsequent ATP molecule detaches myosin, and its hydrolysis allows the head to recover for the next cycle (Lymn and Taylor, 1971). The number of available myosins governs the force generated (Spudich, 2019), and amount of ATP consumed. Therefore, control of myosin availability is paramount in balancing the energy demands of muscle.

Relatively recently, in a field that has a very long history, a new biochemical state of myosin was identified, first in skeletal (Stewart et al., 2010) and then in cardiac muscle (Hooijman et al., 2011). Using a pulse-chase approach where MANT-ATP was perfused into relaxed permeabilized muscle fibres before rapid dilution with unlabelled ATP, a bi-phasic fluorescence decay was observed by fluorescence microscopy, corresponding to the release of MANT-ADP. The faster phase was in part attributed to the normally relaxed turnover of ATP by myosin (DRX state; not activated by actin binding) and the second slower phase(s) was termed the super-relaxed state (SRX). This state potentially solves the energy paradox by acting as a reserve of myosins to be called upon when needed. It is also possible that a subset of the SRX is represented by a structural folded state termed the interacting heads motif (IHM) however discussion over this is hotly debated and lies outside the scope of this article (for a review see (Craig and Padron, 2022)). Over the subsequent years ubiquitous use of a version of this MANT-ATP assay was employed to identify the presence of the SRX state. However, the assay used a spectrophotometric approach that relied on the change in fluorescence of MANT-ATP upon binding to myosin (Hiratsuka, 1983; Cremo et al., 1990 1990; Woodward et al., 1991 1991; Myburgh et al., 1995). These assays have also been used beyond myofilaments to reveal two phases in stopped-flow measurements of MANT-ADP release for purified/recombinant myosins (Anderson et al., 2018; Gollapudi et al., 2021). However, the use of the MANT-ADP release assay was cast into doubt for studies using purified proteins without adequate secondary measures (Walklate et al., 2022; Mohran et al., 2024). This has led to uncertainty in the field and requires a unifying approach.

We and others developed a new single molecule imaging assay to measure the binding and release of fluorescent Cy3-ATP in the milieu of myofibrils (Nelson et al., 2020; Pilagov et al., 2023). These preparations are ideal for studying myosin activity because they preserve the sarcomeric structure while also being small enough to minimize diffusion times. This latter point is of concern, since rate constants of nucleotide release will be sensitive to permeability. The highly permeable nature of myofibrils has long been established (Stehle et al., 2009) with solution exchange on the sub-ms timescale (Solzin et al., 2007). However, although these single molecule techniques provide very detailed information on the spatial location of myosin activity, they are also very complex and time-consuming. Therefore, a middle ground technique is needed. To bridge this gap, we present an approach here to measure the activity of myosins in myofibrils directly using bulk fluorescence imaging. This technique uses a pulse-chase approach based on that of the Cooke lab (Hooijman et al., 2011), but with the very high permeability of myofibrils, and using Cy3-ATP instead of MANT-ATP.

## Results and discussion

Figure 1a shows an overview of the approach, myofibrils from porcine cardiac left ventricular tissue are immobilised on a coverslip surface and incubated with 1 µM Cy3-ATP (in this study Cy3-ATP is a mixture of 2’ and 3’ attachments to the ribose (Toseland and Webb, 2011)) with 500 µM unlabelled ATP to ensure relaxed conditions for 30 mins before performing a rapid wash step, which clears out the fluorescent ATP and washes in 5 mM unlabelled ATP. Following this step images of each myofibril were taken for 15 ms every 5 seconds, and the laser illumination terminated between images to eliminate photobleaching. During the dark period we move the stage to visualize up to three more myofibrils in the imaging chamber before returning to the first. To achieve this, we built our own proof-of-concept stage system with Arduino controllers to coordinate with laser flashes, however commercial automation would reduce the dead-time between images and increase multiplexing. We continued to take images for 30 minutes and then during post-imaging we select a region of the myofibril that does not include any edges using FIJI (ImageJ, NIH). The fluorescence intensity in this region is projected through time to produce a transient of nucleotide release (Figure 1b). The transient is best fit to either a double or triple exponential decay, determined by examination of the amplitude and systematic noise of the residuals for each fit (Fig 1c). The fastest phase (> 0.041 s^−1^) is likely due to remnants of the Cy3-ATP washout, actin-induced ADP release, and/or non-specific binding of nucleotide (Amrute-Nayak et al., 2014; Ušaj et al., 2021) and so is ignored for the remainder of the analysis. However, the remaining two phases fall into the rate constant regime consistent with the DRX (0.041 > 0.007 s^−1^) and SRX (< 0.007 s^−1^) states of myofibrils (Cooke, 2011; Walklate et al., 2022). Fig 2a shows how these rate boundaries were objectively determined. The rate versus percent population of the two slower phases is plotted for every untreated myofibril studied (filled black circles) and then fitted to a Gaussian mixtures model, which uses a statistical approach to cluster datapoints together. This creates a probability density around the cluster point shown as coloured contours in Fig 2a (yellow at peak, blue towards edges). By defining the boundary at 99.95% between populations (Fig 2a: black contour line) an equal number of rate constants in both the DRX and SRX regimes are seen for the untreated data. The average rate constant for DRX and SRX was 0.021 s^−1^ ± 0.002 and 0.0021 s^−1^ ± 0.0005 (errors are SEM, n=10 myofibrils) respectively using this approach.

**Figure 1:**
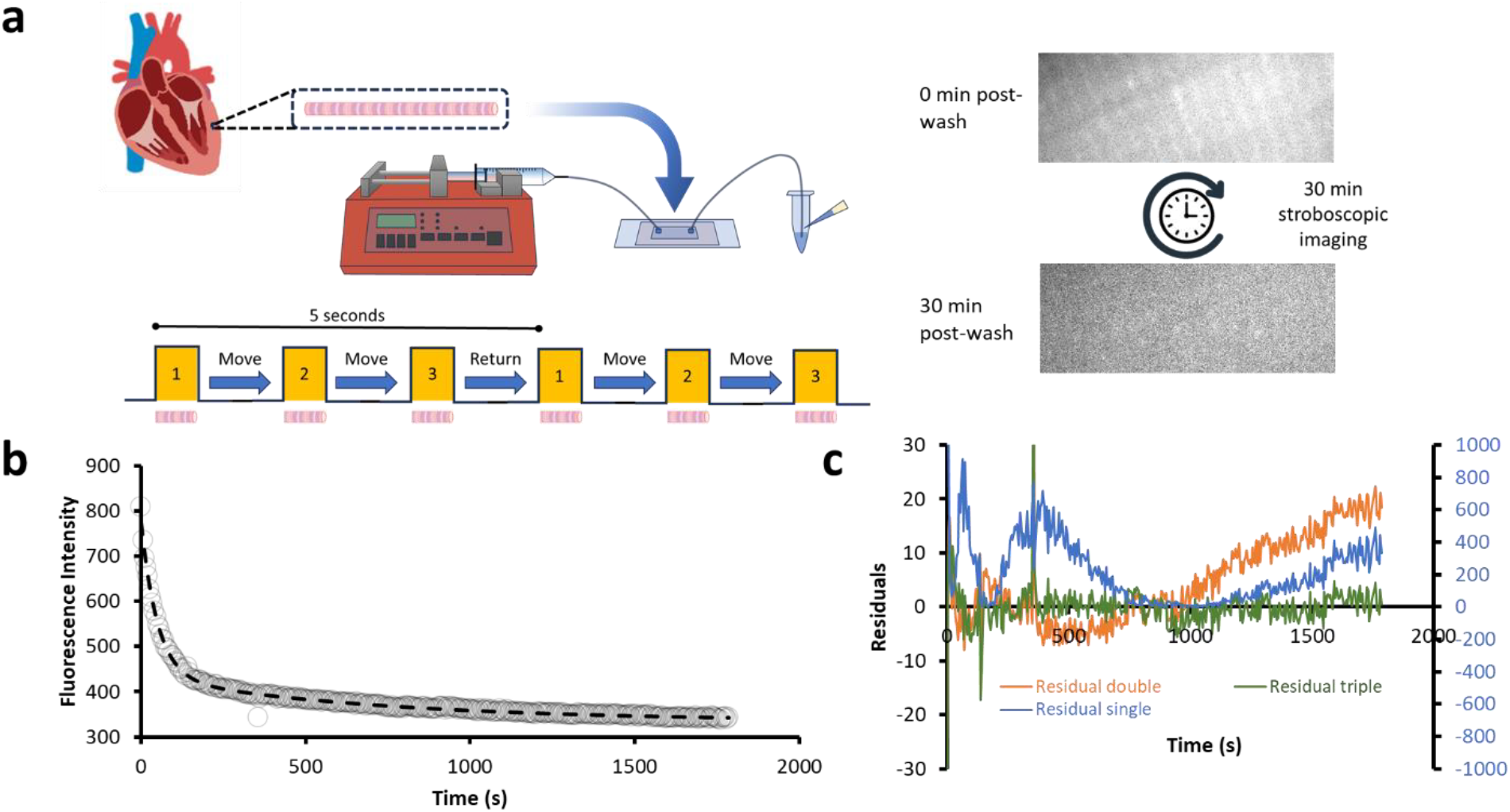
Overview of the method for rapid multiplexed imaging of myofibrils. (**a**) Myofibrils are purified from left ventricular cardiac trabeculae and then injected into a microfluidic cell. After wash-in and then wash-out of Cy3-ATP, the bulk fluorescence over the myofibril is measured and shows a decay over the 30-minute imaging period. (**b**) The fluorescence decay is best fitted as shown to a triple exponential. This is determined by examination of the (**c**) residuals shown for single (blue), double (orange) and triple (green) exponential fits. The single exponential fit residuals are plotted against the right Y-axis showing the large differences from the data. For the double and triple exponentials, these are plotted against the left Y-axis scale. The triple exponential fit (green) is preferred because it shows a better fit throughout the time range particularly at the longer time periods where it is important to obtain a good fit. All data were fit using a logarithmic weighting (due to their exponential distribution), therefore the residuals do not average on zero as would be expected from linear weighting.

**Figure 2:**
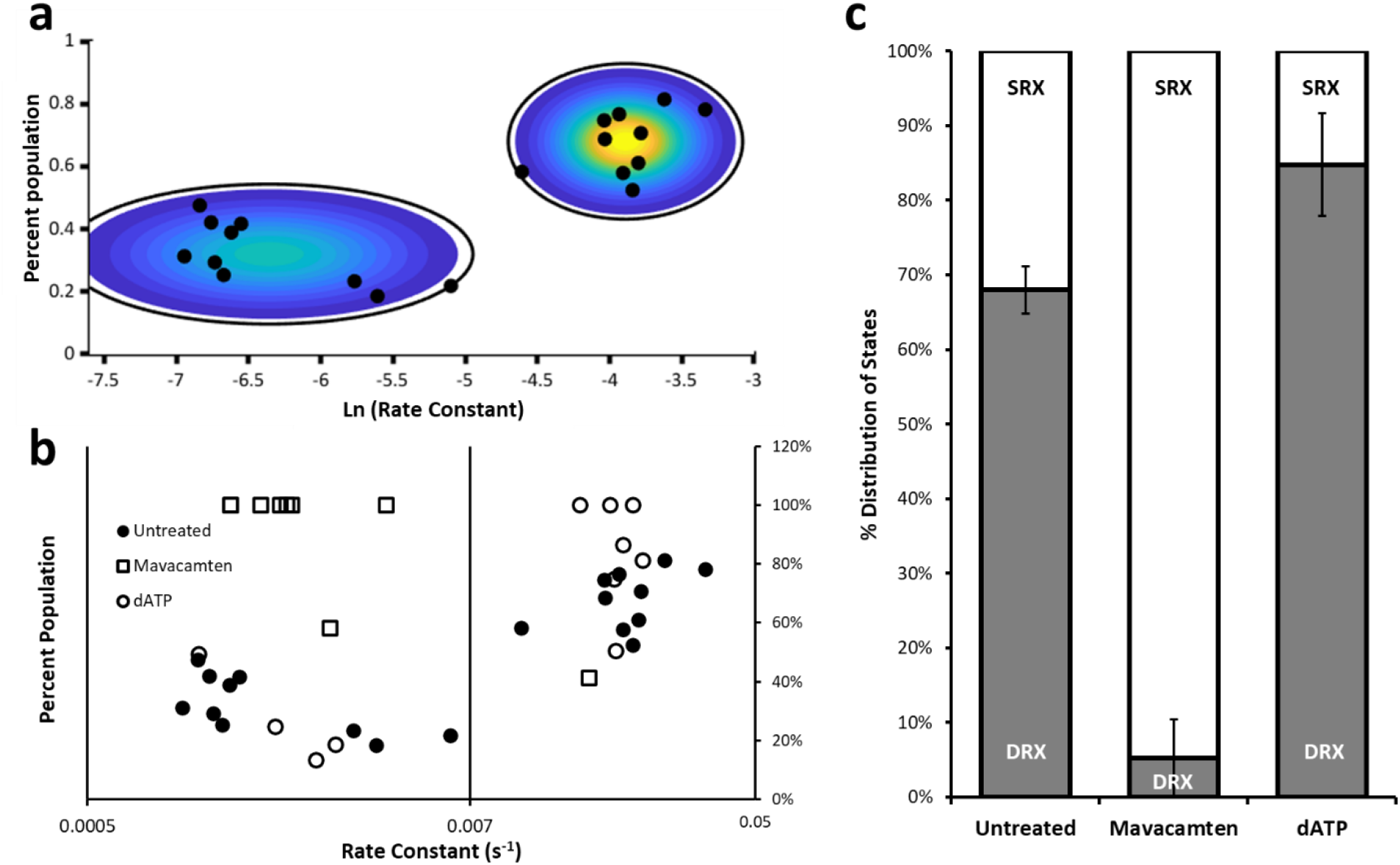
Perturbing the DRX/SRX ratio using Mavacamten and dATP. (**a**) Untreated data fitted to a Gaussian mixtures model shows two major populations. The colour distribution reflects the probability density of the 2D Gaussian used for fitting and the surrounding black circle is the density at 99.95% probability; this was used to independently define the boundaries between the populations for the SRX vs DRX populations. (**b**) Rate constants for all myofibrils are plotted versus their percent amplitude (taking only the slower two phases of the transients (see Figure 1b)). Values with only a single population are marked at 100%. This representation provides an overview of all the data, showing untreated (filled circles), treated with 30 µM Mavacamten (squares) and 100% dATP (open circles). The vertical line in the graph represents the boundary rate constant defined by the Gaussian mixtures model. For clarity, (a) and (b) are aligned. (**c**) Provides a percentage of the DRX/SRX populations determined from the fitted data for each myofibril averaged across all measured myofibrils. This chart shows relative to DRX in untreated 68.0% (± 3.2 SEM n = 10 myofibrils) there is a significant decrease (p<0.002; Student’s T-test) in the presence of mavacamten to 5.2% (± 5.2 SEM n = 8 myofibrils). With deoxyATP (dATP), the total amount of DRX is significantly increased (p<0.05; Student’s T-test) to 94.8% (± 8.0 SEM n = 7 myofibrils).

To test that the two rate constants reflect the two populations of SRX and DRX we perturbed the DRX/SRX ratio by washing out the Cy3-ATP with a solution containing 1) 30 µM mavacamten + 5mM unlabelled ATP or 2) 100% deoxyATP. These were chosen because they provide the expected lower (mavacamten) and upper (dATP) limits on the amount of DRX. Mavacamten has been shown to reduce activity in purified myosin (Anderson et al., 2018), myofibrils (Walklate et al., 2022), and patients (Zampieri et al., 2021). It is the only myotropic drug to receive FDA-approval for the treatment of the cardiac condition obstructive hypertrophic cardiomyopathy. Conversely, dATP has been shown extensively to activate purified myosin, muscle fibres and myofibrils (Regnier et al., 2000; Walklate et al., 2022; Mohran et al., 2024). In many of the cases with these myotropes, we observed only one of the two slower phases (DRX or SRX) indicating a full shift in distribution of the population for that myofibril. Plotting the observed populations as we did for untreated revealed again two clear populations for dATP treated myofibrils (Fig 2b open circles) with average rate constant per regime DRX = 0.020 s^−1^ (± 0.001 SEM n = 7 observations), and SRX = 0.002 s^−1^ (± 0.003 SEM n = 4 observations)), three of the seven measured samples had only DRX with no apparent SRX population (plotted as 1--% DRX). Overall, this resulted in an elevated percentage population in DRX relative to untreated (Fig 2c). By contrast, mavacamten treatment showed only a single instance of a myofibril with a rate constant that fell into the DRX category (rate constant = 0.016 s^−1^). The remainder were entirely SRX leading to a large overall shift towards SRX (Fig 2c) with an average rate constant = 0.002 s^−1^ (± 0.0003 SEM n = 8 observations). This is expected at the saturating concentrations of mavacamten used and does not indicate if there would be intermediate rate populations as suggested from stopped-flow experiments (Mohran et al., 2024). To quantitate the amount of DRX, we determined the amplitudes of the transients per myofibril and then plotted these as an average across all myofibrils studied (Fig 2c). These data show that untreated porcine myofibrils possess 68.0% DRX, whereas upon treatment with mavacamten, the DRX population drops significantly to 5.2% and with dATP there was an increase in DRX to 84.8% (errors in figure 2 legend). These results are in close agreement with previously published data for the effects on porcine myofibrils (Walklate et al., 2022; Pilagov et al., 2024), indicating this method provides straightforward analysis across the full range of DRX proportions measured to date.

## Conclusions

We show here that it is possible to image the fluorescence decay following a rapid washout of Cy3-ATP to detect at least two phases in cardiac myofibrils. This approach builds on the pioneering work from the Cooke lab and offers a fast, reliable way to make measurements of the cardiac myosin reserve within the myofilament lattice. Cy3-ATP provides a strong fluorescence signal that does not affect the activity of myosin (Conibear et al., 1996; Amrute-Nayak et al., 2014; Ušaj et al., 2021), and the use of myofibrils ensures the myofilament lattice structure but with the advantage rapid diffusion necessary for chase experiments. Furthermore, our method utilizes rapid flashes of the excitation laser with a dark period between frames, enabling the measurement of multiple myofibrils in one imaging chamber. Because it takes 30 mins to accurately measure the proportion of SRX, multiplexing offers a means to accelerate data collection, rather than observing myofibrils in series. Here we show that mavacamten nearly eliminates the DRX phase and that deoxyATP reduces the contribution of the SRX phase, providing the upper and lower bounds of the response. We have presented this technique to enable future studies using any myofibril source, including those containing mutations isolated from patients or stem cell derived, or to test for other compounds that alter the DRX/SRX ratio. Such studies may begin to shed light on perhaps the most pressing question, which is how the DRX/SRX ratio is maintained and modulated. Rapid interconversion between these populations will lead to a single phase (Walklate et al., 2022; Mohran et al., 2024), however, our measurements show two phases, and their rate constants are maintained despite drug treatment indicating that these populations likely do not interconvert in relaxed muscle.

Although we have demonstrated a simple, rapid tool to measure and quantify myosin activity in a near-native environment, other methods offer their own valuable insights such as polarization, FRET, SHG and X-ray diffraction that measure the physical positions of the heads (Nucciotti et al., 2010; Kampourakis and Irving, 2015; Fusi et al., 2016; Irving and Craig, 2019; Chu et al., 2021; Ma et al., 2022; Rasicci et al., 2022; Brunello and Fusi, 2024), which may relate to myosin’s biochemical states. Nonetheless, directly imaging fluorescent ATP release from myofibrils adds significantly to the tools that can be used to fully solve the origin of the paradox of energy usage in the heart and other muscle tissues.

## Methods

No live animals were used in this study. Muscle tissue was collected in accordance with the U.K. Animals (Scientific Procedures) Act 1986 and associated guidelines.

### Dissection and storage of porcine heart

The heart of a freshly euthanised adult farm pig was submerged in ice-cold cardioplegic solution (5.5 mM Glucose, 0.5 mM MgSO_4_, 24 mM KCl, 20 mM NaHCO_3_, 109 mM NaCl, 0.9 mM H_2_NaO_4_P, 1.8 mM CaCl_2_, 0.01% (w/v) NaN_3_, pH 7.4) immediately after excision. Whilst submerged, left ventricular trabecular samples were dissected and, cut into ~5 mm thick pieces, individually wrapped in foil, immediately flash frozen and stored long-term at −80 °C.

### Myofibril isolation

Porcine left ventricular trabecular myofibril suspensions were prepared as follows: a flash-frozen sample of porcine left ventricular trabeculae was rapidly thawed in chilled Prep buffer (20 mM MOPS, 132 mM NaCl, 5 mM KCl, 4 mM MgCl_2_, 5 mM EGTA, 0.01% (w/v) NaN_3_, 5 mM DTT, protease inhibitor cocktail (A32965; Thermo Scientific), pH 7.1 at RT) (Vikhorev, Ferenczi, and Marston 2016) and cut into ~1 mm thick strips. Sample strips were transferred to 500 µl of Prep buffer in a 2 ml microcentrifuge tube for homogenization using a Tissue Ruptor II (Qiagen) at medium-low speed for 10 seconds twice, with a 1-minute rest on ice between. Homogenized tissue was then permeabilized for 30 minutes at 4 °C, rotating, in Permeabilization buffer (Prep buffer + 1% Triton X-100). Myofibrils were collected by centrifugation at 1000 g for 3 minutes and resuspended in chilled Prep buffer. Washes were repeated 3 times to remove all traces of Triton X-100 and myofibril suspensions were diluted or concentrated where necessary to obtain an OD_600_ of ~0.6. Myofibril suspensions were stored at 4 °C and discarded at the end of each day.

### Microfluidic flowcells

Microfluidic flowcells were constructed by adhering a plasma-cleaned borosilicate coverslip (Menzel Gläser, 24 × 40 mm, 1.5 thickness), coated in 5 µg/ml >300KDa poly-L-lysine (PLL, Sigma) to a standard microscope slide prepared for microfluidics, using a 360 µm thick adhesive gasket. Microscope slides were prepared for microfluidics by drilling two 1.5 mm diameter holes, 1.5 cm apart and adhering polyethylene tubing (Gradko, GE-0086-033, 0.86 mm internal diameter, 0.33 mm wall) to each using an acrylic bonder (RS, 144-406). Flowcells were connected to a syringe pump (WPI, AL-1000) at one end and at the other, to a 1.5 ml microcentrifuge tube with holes to allow access of pipette tips. 200 µl of myofibril suspension was added to the 1.5 ml microcentrifuge tube and drawn into the flowcell at 1 ml/min then washed back through at 0.2 ml/min. To allow adhesion of myofibrils to the PLL-coated surface, flowcells were incubated on ice for 30 minutes. Excess non-adherent myofibrils were withdrawn from the flowcell using 600 µl Prep buffer at 10 ml/min.

### Image acquisition

All imaging was carried out at room temperature (21°C) using a custom-built oblique angle fluorescent (OAF) microscope (Desai, Geeves and Kad 2015). The sample was illuminated for 15 ms every 5 seconds using a 561 nm laser (OBIS LS laser, Coherent, USA) to excite Cy3-ATP, for a total time of 30 mins.

Myofibrils were incubated with Cy3-ATP buffer (Prep buffer + 1 µM Cy3-ATP, 500 µM ATP, 0.5 mM phosphoenolpyruvate (PEP), 2.2 units Pyruvate kinase (PK)) for 30 mins at RT. Following this, a single frame was taken of the myofibril fully saturated with Cy3-ATP, followed by a rapid wash with Chase buffer (Prep buffer + 5 mM ATP, 5 mM PEP, 22 units PK) at 10 ml/min. To increase efficiency, flowcells were mounted onto a custom-built automated stage programmed to move between user-specified positions along one axis. Between each frame images from neighbouring myofibrils were taken. Use of this automated stage allowed up to 4 myofibrils to be imaged during the 30-minute imaging period.

Mavacamten (MedChemExpress) was solubilized in 100% DMSO and added to the Chase buffer at final concentration of 30 µM with a final DMSO of 0.3% v/v. For dATP experiments, 5 mM dATP (NEB N0440S) was used in the Chase buffer replacing unlabelled ATP.

### Data analysis

To extract the rate of fluorescence decay for each myofibril, an ROI was drawn using ImageJ to encompass the entire myofibril and the total intensity profile was extracted through time. These data were transferred to Excel and then fitted to the exponential decays as shown in figure 1 to provide both rate constants and amplitudes.

For the Gaussian mixture model we used Matlab’s in-built function with diagonal covariance and the number of components fixed at two. The mixtures calculation was iterated 1000 times for best fit and then repeated 100 times to determine the distribution of best fit means and standard deviations for DRX and SRX. These values were then used to calculate the boundaries between DRX and SRX by altering the multiple of standard deviations that led to an equal population of DRX and SRX.

